# Environmental gradients shape climate adaptive phenotypes in Muscovy ducks in Benin with implications for adaptation policy

**DOI:** 10.64898/2026.06.05.730266

**Authors:** Lionel Kinkpe, Christian O.V. Chabi, Solomon I. Ahamba, Zeleke Tesema, Naqash Goswami, Niu Yurui, Rauan Abdessan, Daouda Libanio, Camus M. Adoligbe, HU Zhigang, Wang Xia

## Abstract

Climate change threatens livestock-dependent livelihoods across sub-Saharan Africa, yet National Adaptation Programmes of Action (NAPAs) and Nationally Determined Contributions (NDCs) rarely incorporate animal genetic resource management despite explicit mandates under the Convention on Biological Diversity’s Global Plan of Action for Animal Genetic Resources. This study addresses this policy gap by investigating how environmental gradients shape phenotypic adaptation in Muscovy duck (*Cairina moschata*) populations across Benin’s three agroecological zones, generating evidence to inform climate-responsive livestock policy. We surveyed 505 farmers and characterized 1,300 ducks across dry (Temperature-Humidity Index [THI]: 70-74), sub-humid (THI: 74-78), and humid (THI: 78-82) zones. Climate analysis confirmed accelerating thermal stress (+0.38 degrees C per decade in humid zones), exceeding IPCC West African projections and threatening Sustainable Development Goal 2 (Zero Hunger) targets. Principal component analysis revealed zone-specific phenotypic clustering (57.9% variance explained), with thermoregulatory traits showing coordinated adaptation: body volume-surface ratios 42.7% lower in humid zones (p = 0.002), limb proportions 72.7% higher (p < 0.001), and white feather coverage 31.3% greater (p = 0.019). Reproductive performance declined significantly along the thermal gradient (lifetime production: 40.7 vs. 37.1 eggs; p < 0.001), quantifying climate-driven productivity losses relevant to national food security assessments. Phenotype-climate associations demonstrated that locally adapted variants (black-white pied in dry zones; white-black-green pied in humid zones) maintain superior fitness under native conditions, providing critical evidence for the breed-for-environment approach advocated by FAO’s Climate-Smart Agriculture framework. These findings provide an empirical foundation for incorporating animal genetic resource conservation into Benin’s NDC implementation, ECOWAS regional livestock strategies, and the African Union’s Comprehensive Africa Agriculture Development Programme (CAADP) climate resilience targets.

## 1. Introduction

Climate change represents a growing and systemic threat to livestock-dependent livelihoods across sub-Saharan Africa, where hundreds of millions of people rely on animal agriculture for food security, income generation, and cultural identity [1, 2]. Livestock systems in West Africa are particularly exposed to rising temperatures, altered rainfall patterns, and increasing climate variability. The IPCC Sixth Assessment Report (AR6) indicates that under intermediate emission pathways, regional warming across West Africa is projected to exceed 1.5 °C by mid-century, with higher levels of warming possible later in the century, accompanied by increased heat stress, changing disease dynamics, and constraints on feed and water availability [3].

These climatic pressures have important implications for poverty reduction and food security. According to the World Bank and assessments synthesized by IPCC AR6, climate change could push between 32 and 132 million additional people into extreme poverty globally by 2030, with Africa among the most affected regions due to high climate sensitivity and dependence on climate-exposed livelihoods [3, 4]. Such trends pose significant risks to progress toward Sustainable Development Goal 2 (Zero Hunger) and SDG 1 (No Poverty), underscoring the urgency of adaptation strategies that strengthen the resilience of livestock-based food systems.

Despite this urgency, animal genetic resource management remains weakly operationalized within national climate adaptation frameworks across much of sub-Saharan Africa. While National Adaptation Programmes of Action (NAPAs) and Nationally Determined Contributions (NDCs) frequently emphasize agricultural resilience, they often provide limited concrete guidance or investment pathways for the conservation and adaptive use of livestock genetic diversity [5]. This gap persists despite longstanding international commitments. The Global Plan of Action for Animal Genetic Resources, adopted under the auspices of the FAO, explicitly identifies the characterization, conservation, and sustainable use of livestock genetic diversity as a cornerstone of adaptation to environmental change [6]. Similarly, the Convention on Biological Diversity recognizes the safeguarding of genetic diversity in domesticated animals under Aichi Biodiversity Target 13, and the Paris Agreement frames adaptation as a global challenge requiring integrated, ecosystem-based and knowledge-inclusive approaches [7]. However, translation of these global commitments into actionable livestock-sector policies at the national level remains inconsistent, creating a persistent disconnect between biodiversity obligations and practical climate adaptation planning in animal agriculture.

Benin exemplifies a broader and well-documented challenge across sub-Saharan Africa: the limited translation of climate adaptation objectives into operational livestock genetic resource–focused strategies. Comparative analyses of livestock adaptation pathways across Africa consistently show that, although livestock systems are widely recognized as highly climate-vulnerable, explicit integration of animal genetic diversity, breed characterization, and genotype–environment matching remains weak in practice [5, 8]. Across West African contexts, adaptation planning has largely emphasized productivity enhancement through feed, health, and management interventions, while comparatively little empirical attention has been directed toward conserving and deploying locally adapted genetic resources [9]. As a result, policymakers often lack a robust scientific evidence base linking specific livestock populations to climate resilience outcomes, constraining the development of targeted, genetics-informed adaptation strategies.

Environmental gradients provide a powerful and widely accepted framework for addressing this evidence gap, as they enable inference of adaptive trait–environment relationships without requiring long-term temporal datasets. In evolutionary biology and ecological genomics, gradient-based approaches have long been used to identify local adaptation, fitness trade-offs, and adaptive constraints across heterogeneous environments [10]. More recently, these approaches have been increasingly applied in livestock systems to identify thermotolerant phenotypes and climate-adaptive genetic variation under real production conditions, particularly in low-input systems exposed to climatic stressors [11, 12].

Benin represents a particularly suitable setting for such analyses due to its pronounced north–south environmental heterogeneity, spanning arid and semi-arid Sudanian zones to humid Guinean ecosystems over a relatively short geographic distance. Agro-ecological studies have documented strong spatial variation in rainfall regimes, ambient temperature, and humidity across this gradient, with direct implications for livestock heat stress exposure and productivity [13, 14]. In poultry research, thermal stress is commonly quantified using the Temperature–Humidity Index (THI), a physiologically meaningful metric shown to be closely associated with growth performance, immune competence, and survivability under tropical conditions [15, 16]. Spatial variation in THI across West African agro-ecological zones therefore constitutes a natural experimental framework for identifying heat-tolerant phenotypes that may become increasingly valuable under ongoing warming.

Within this context, Muscovy ducks (Cairina moschata) represent a strategically important yet under-studied genetic resource for climate adaptation in tropical Africa. Empirical population-level studies demonstrate substantial phenotypic diversity in African Muscovy ducks shaped by long-term exposure to diverse agro-ecological conditions under extensive management systems [17, 18]. Compared with commercial poultry lines, Muscovy ducks exhibit greater tolerance to heat stress, nutritional variability, and endemic diseases under smallholder conditions [19]. Despite these attributes, Muscovy ducks remain largely absent from quantitative climate adaptation research, underscoring the need for integrative studies that explicitly link phenotypic diversity, environmental gradients, and adaptive potential.

The limited integration of locally adapted livestock genetic resources into climate adaptation planning carries important consequences for environmental management and system resilience. In the absence of systematic phenotypic and performance characterization, locally adapted Muscovy duck populations are vulnerable to progressive genetic erosion driven by unmanaged crossbreeding and replacement by exotic breeds promoted through development interventions [20]. Such processes have been widely documented across livestock species in sub-Saharan Africa and are associated with the loss of adaptive traits critical for coping with heat stress, disease pressure, and resource variability under climate change [21]. This erosion reduces the adaptive capacity of livestock systems precisely as environmental stressors intensify.

Equally important, the lack of quantitative performance data collected across differentiated environmental conditions constrains the ability of decision-makers to formulate evidence-based recommendations for strategic breed deployment. Recent syntheses emphasize that adaptation planning in livestock systems continues to rely heavily on generalized productivity and management interventions, while underutilizing the potential of locally adapted genetic resources due to insufficient empirical evidence linking phenotype to environmental performance [5, 8].

This study addresses these gaps by examining phenotypic diversity and reproductive performance of Muscovy ducks across Benin’s environmental gradient. By integrating climatic variation, morphometric traits, reproductive outcomes, and production systems, the analysis generates spatially explicit evidence linking livestock genetic resources to climate adaptation potential. Such evidence is essential for informing environmental assessment frameworks that seek to conserve adaptive diversity and strengthen the resilience of smallholder livestock systems under increasing climatic uncertainty.

## 2. Materials and Methods

### 2.1. Ethics statement

This study involved non-invasive phenotypic characterization of privately owned Muscovy ducks under normal husbandry conditions. No experimental treatments, invasive procedures, blood sampling, or tissue collection were performed.

According to national regulations in Benin, formal institutional animal ethics approval is not required for observational field studies involving routine morphometric measurements conducted with owner consent.

All procedures were conducted with the informed consent of animal owners and in accordance with internationally recognized animal welfare standards.

### 2.2. Study Area: Policy-Relevant Environmental Stratification

The study was conducted across Benin (6°10’–12°25’N, 0°45’–3°55’E) and stratified according to the three agroecological zones recognized in national agricultural planning frameworks (Figure 1). Figure 1 illustrates the spatial distribution of the study areas across these agroecological zones, differentiated by long-term (40-year) gradients of temperature, relative humidity, and annual precipitation. This zonation, established by Benin’s Institut National des Recherches Agricoles du Benin (INRAB), ensures findings are directly applicable to existing policy structures. The dry Sudanian zone corresponds to Borgou and Alibori departments (national priority areas for livestock development); the sub-humid transition zone encompasses Collines, Zou, and Plateau departments; and the humid Guinean zone includes Oueme, Mono, and Atlantique departments. This tri-zonal framework captures the north-south precipitation gradient (900-1,400 mm annually), temperature gradients, and the ecological transitions that structure agricultural potential across Benin [22].

**Figure 1.**
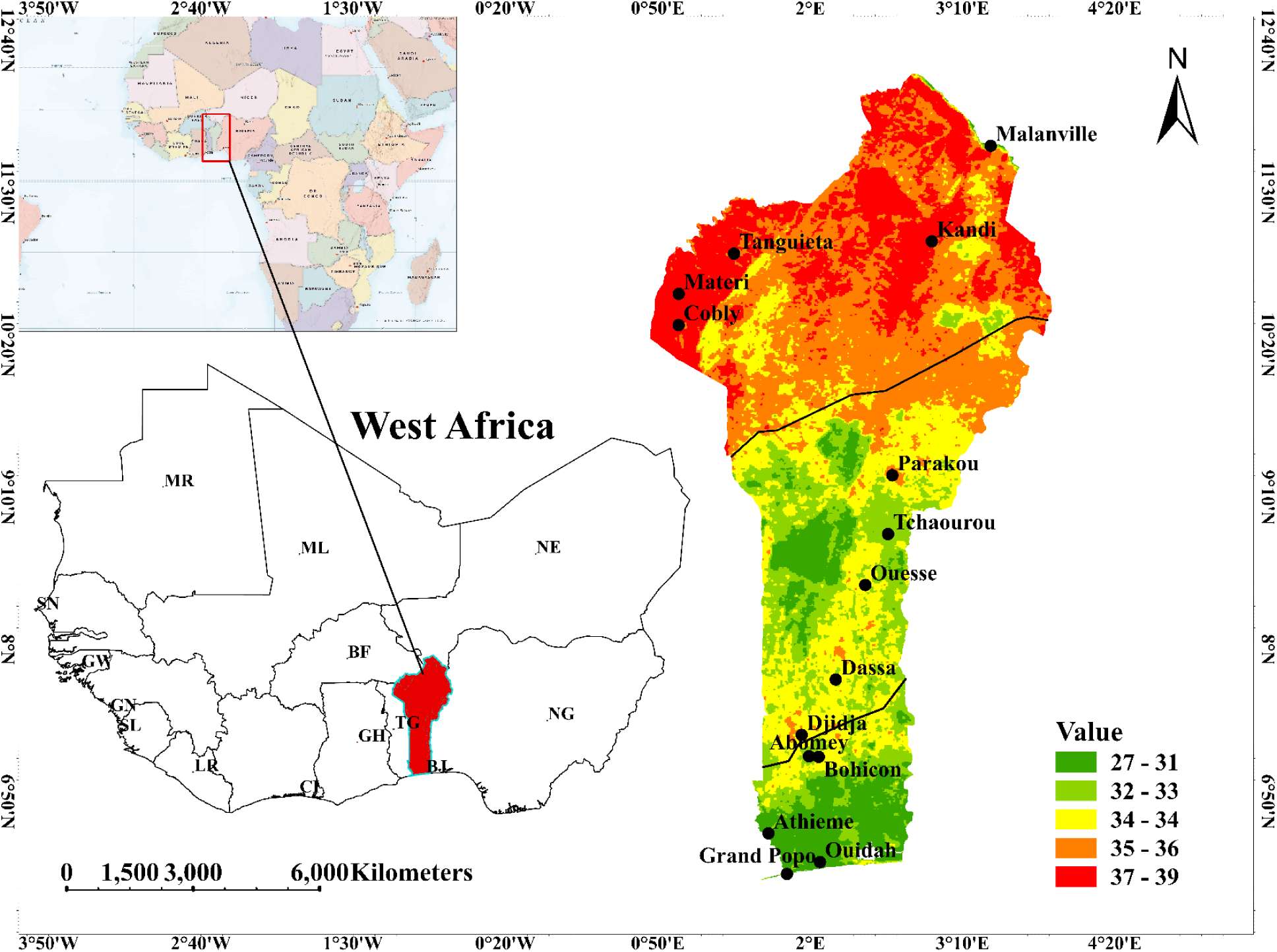
Spatial distribution of study areas across Benin’s agroecological zones based on long-term climatic conditions. The map illustrates the geographic location of the study areas and their stratification into three agroecological zones defined by long-term (40-year) averages of temperature, relative humidity, and annual precipitation, reflecting the north–south climatic gradient that structures agricultural and livestock production systems in Benin.

Long-term climate records (1979-2022) were obtained from Tutiempo.net for survey locations, including monthly maximum (Tx), minimum (Tn), and mean (Tmean) temperatures, as well as total monthly precipitation, and relative humidity. These data enable trend analysis relevant to national climate vulnerability assessments conducted under UNFCCC reporting requirements. The Temperature-Humidity Index (THI) was calculated following [23]:

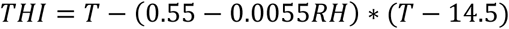

where T is annual average temperature and RH is relative humidity and RH is relative humidity (%).

Temperature–humidity index (THI) thresholds for poultry heat stress onset (THI ≥ 70), severe stress (THI ≥ 78), and critical stress (THI ≥ 82) were applied following established international poultry heat-stress classifications [24].

While long-term climate records (1979–2022) were used to characterize agroecological gradients and to demonstrate the magnitude and direction of climate change across Benin, phenotypic and survey data were necessarily limited to the years of field investigation. Historical phenotypic records spanning the full 40-year period are not available at the national scale for Muscovy duck populations. Consequently, phenotypic characterization and farmer surveys reflect current populations sampled across the country, and climate variables corresponding to the study years and locations were used for phenotype-environment association analyses. The long-term climate dataset therefore served to contextualize environmental change and policy relevance, whereas phenotype-climate relationships were evaluated using contemporaneous climate conditions representative of the survey period.

### 2.3. Sampling Design and Stakeholder Engagement

In the absence of a livestock census including duck populations, a data gap identified in Benin’s NAPA as limiting adaptation planning, a stratified non-probability sampling approach was employed. Initial participant identification was facilitated by Techniciens Specialises en Production Animale (TSPA), the government livestock extension agents operating under the Ministry of Agriculture. This engagement ensured policy relevance by working within existing institutional structures and created channels for disseminating findings to farmers.

Eligibility criteria targeted experienced producers (at least 2 years duck farming; at least 15 ducks owned) capable of providing reliable performance data. From initial contacts, snowball sampling expanded the sample within each zone. The final sample (n = 505 farmers: dry zone = 119; sub-humid = 284; humid = 102) exceeded Cochran’s minimum threshold (n = 384) for 95% confidence intervals with 5% margin of error. Multiple entry points in each zone and cross-validation of response patterns minimized selection bias. This approach is consistent with FAO guidelines for field-based genetic resource characterization when formal registries are unavailable (FAO, 2012).

### 2.4. Phenotypic Characterization Protocol

Following FAO’s standardized protocols for animal genetic resource assessment (FAO, 2012), a total of 1,300 mature Muscovy ducks were characterized. These protocols are the internationally recognized methodology for generating data compatible with the Domestic Animal Diversity Information System (DAD-IS) maintained under CBD commitments. To ensure consistency, all measurements were conducted by a single trained individual. Quantitative traits measured included body length, sternum length, tarsus length, drumstick length, beak dimensions (length, width), thoracic perimeter, and live weight. Qualitative traits assessed included plumage color patterns and caruncle characteristics.

Derived thermoregulatory indices were calculated based on established avian physiological principles [25, 26]: body volume-surface ratio (thoracic perimeter divided by live weight), beak heat dissipation ratio (beak length divided by width), limb proportion ratio (tarsus length divided by drumstick length), and white feather proportion (white feather coverage divided by body length). These indices operationalize Allen’s and Bergmann’s ecogeographic rules for quantitative assessment of thermoregulatory adaptation.

### 2.5. Statistical analysis of data

The data collected were systematically organized, cleaned, and processed to ensure accuracy and usability before analysis, which was conducted using R version 4.1.0 [27]. Descriptive statistics were used to characterize sociodemographic and management variables. Categorical associations were assessed using Chi-square tests. Generalized linear mixed models (GLMMs) with Poisson distribution (lme4 package) modeled egg production, with agroecological zone as fixed effect and village-level random intercepts (ICC = 0.12, 95% CI: 0.06-0.18). Beta regression (betareg package) modeled proportional outcomes (hatchability, mortality). Principal component analysis (PCA) characterized multivariate phenotypic structure. Between-zone differences in thermoregulatory indices were tested using ANOVA with Tukey’s HSD post-hoc comparisons. Statistical significance was set at alpha equals 0.05.

## 3. Results

### 3.1. Climate trends across agroecological zones

Climate analysis revealed warming trends across all agroecological zones that exceed regional projections reported in the IPCC Sixth Assessment Report (Figure 2). Since 2000, mean temperatures have increased at rates of +0.38 °C per decade in the humid zone, +0.35 °C per decade in the sub-humid zone, and +0.31 °C per decade in the dry zone. These rates are higher than the IPCC AR6 median projection of +0.25 °C per decade for West Africa under the SSP2–4.5 scenario, indicating accelerated warming across the study area.

**Figure 2.**
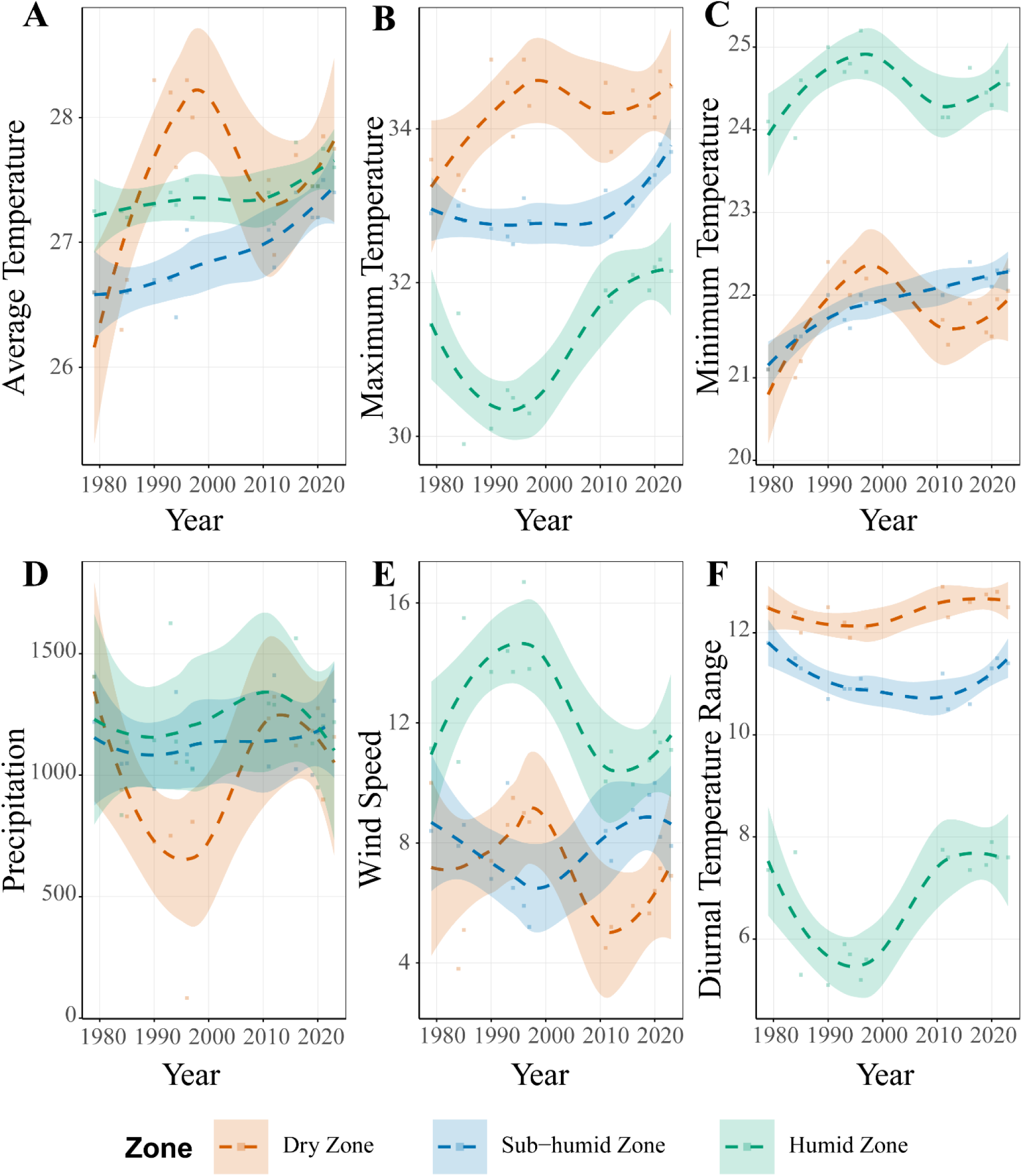
Long-term climate trends across Benin’s agroecological zones (1979–2023). The figure presents locally estimated scatterplot smoothing (LOESS) trends with 95% confidence intervals illustrating non-linear changes in key climatic variables across agroecological zones: (A) average temperature, showing long-term warming across all zones with the steepest increase in the humid zone; (B) maximum temperature, highlighting differential warming patterns in peak daytime temperatures; (C) minimum temperature, indicating nighttime warming and a reduced diurnal temperature range in humid zones; (D) precipitation, characterized by increasing rainfall variability with zone-specific patterns; (E) wind speed, displaying divergent trends with increasing wind speeds in dry zones and decreasing trends in humid zones; and (F) diurnal temperature range, remaining relatively stable in dry zones but exhibiting increased variability in humid zones.

Minimum temperatures increased particularly rapidly in the humid (+0.41 °C per decade) and sub-humid (+0.35 °C per decade) zones, suggesting reduced nocturnal cooling that may intensify cumulative thermal stress. The dry zone exhibited the highest absolute temperatures, with a mean maximum of 38.2 ± 1.3 °C, and the greatest diurnal temperature range (12.5 ± 0.8 °C). Precipitation increased in the humid and sub-humid zones after 2000, while remaining relatively stable in the dry zone.

THI values delineated a clear physiological stress gradient. The humid zone consistently experienced THI values between 78 and 82, exceeding the threshold for severe heat stress, whereas the sub-humid zone (THI 74–78) was characterized by chronic moderate stress. The dry zone exhibited marginal stress conditions (THI 70–74) with high interannual variability. Under continued warming, THI values in the sub-humid zone are projected to reach the severe stress range within the next 15–20 years.

### 3.2 Production Systems: Adaptation Practices and Policy Leverage Points

Survey results revealed that climate-responsive production systems incorporating indigenous adaptation knowledge could inform policy design.

Sociodemographic patterns indicated differential development of duck production across zones (Table1). Duck farming was predominantly managed by men (84.6%), although female participation was highest in the humid zone, a factor relevant for gender-responsive agricultural programming under SDG 5. The majority of farmers (90%) lacked formal training, highlighting opportunities for extension service enhancement. Farming experience averaged 11.5 years in the dry zone compare to 4-7 years elsewhere, suggesting that duck production has been established in drier regions.

Breeding objectives showed clear climate-zone associations with implications for genetic improvement policy (Table 2; Supplementation file Figure S1). Humid zone farmers prioritized dual-purpose production (56.9%), sub-humid zone farmers emphasized conservation (44.7%), while dry zone farmers focused on egg production (44.5%) and conservation (42.9%). These divergent objectives indicate that uniform breed improvement programs would be inappropriate; zone-specific approaches aligned with farmer priorities are essential for adoption.

**Table 1:**
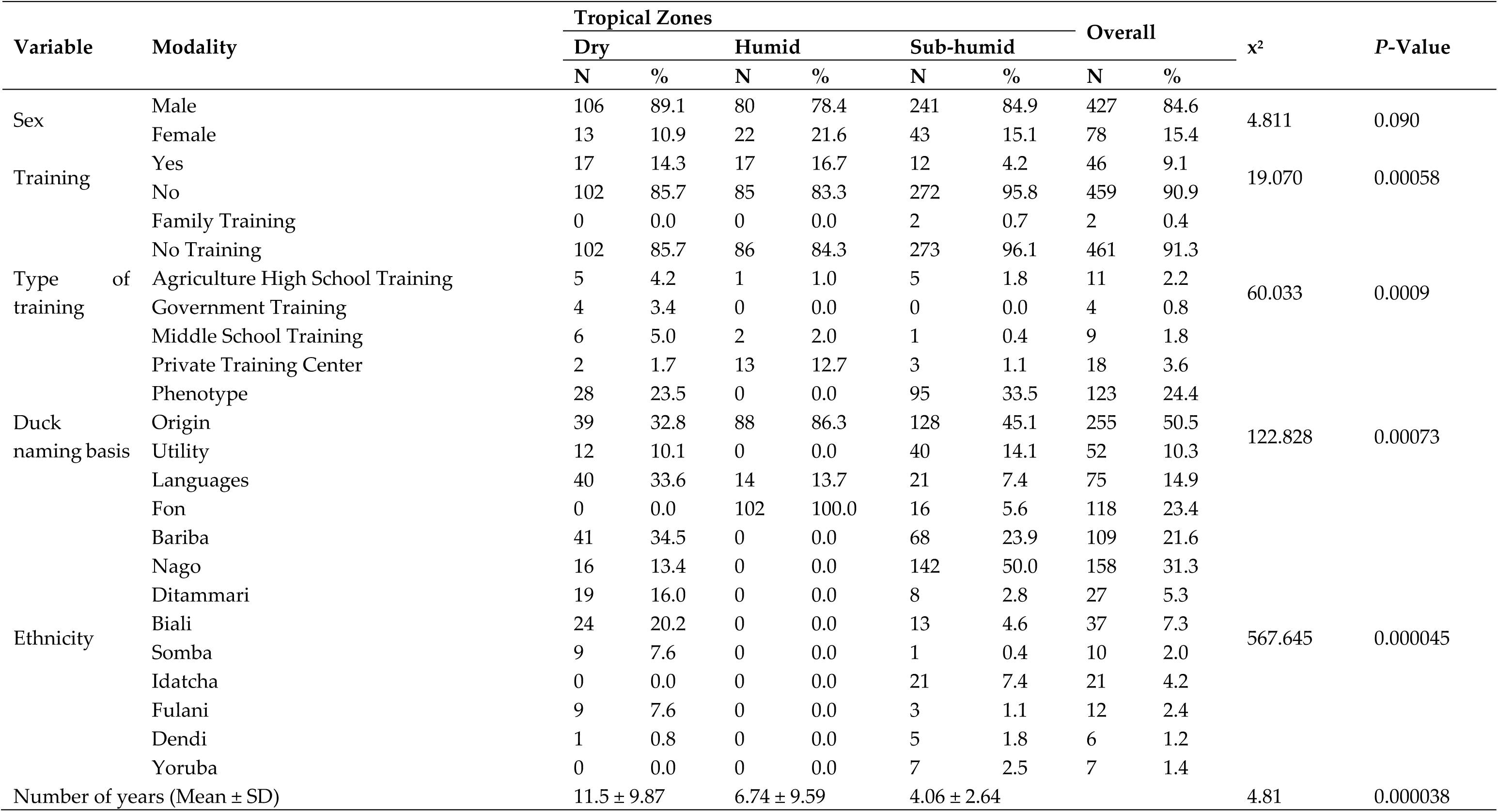
Socio-demographic characteristics of duck farmers in Benin.

**Table 2:**
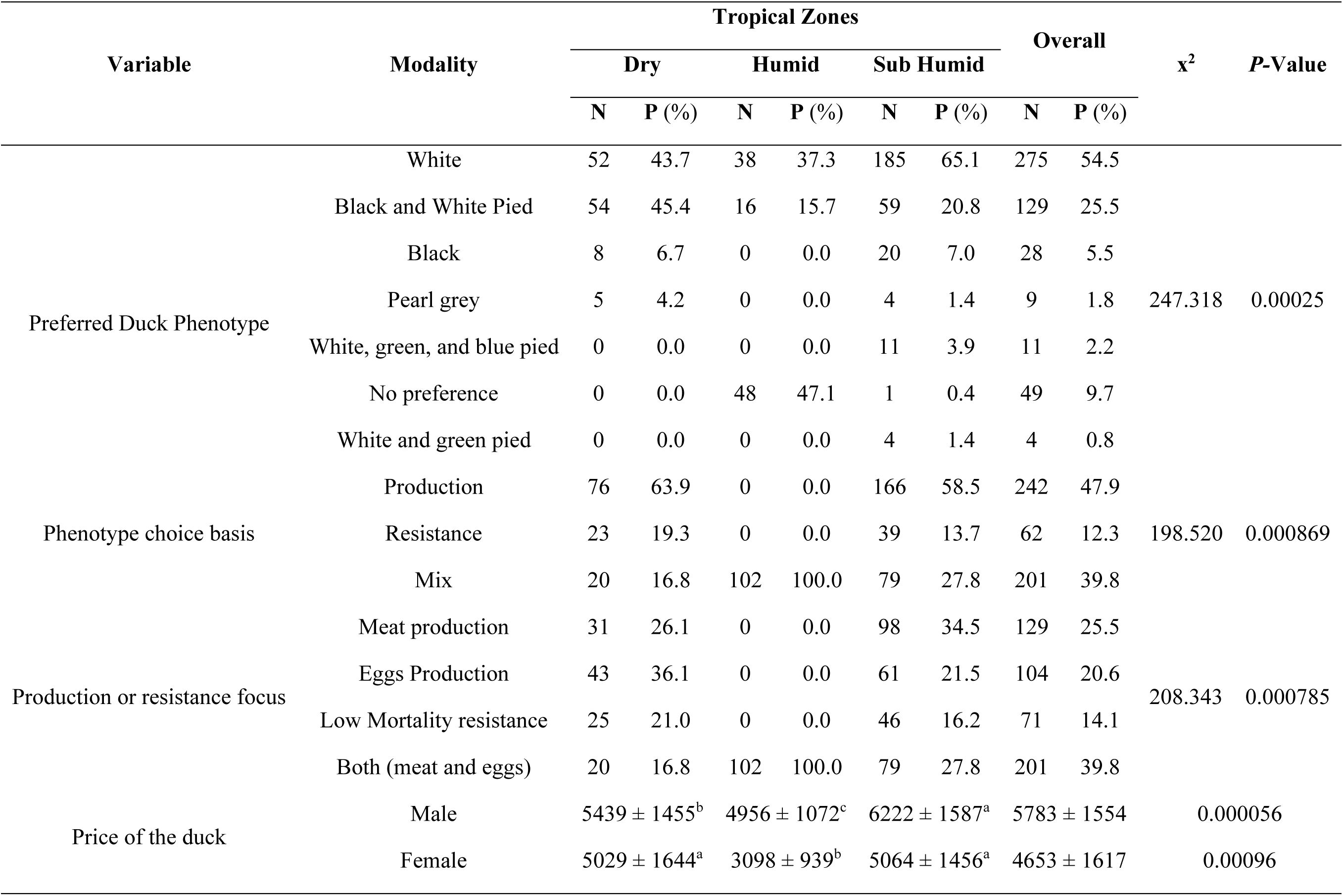
Duck phenotype preferences across Agro-ecological zones.

Management systems demonstrated autonomous adaptation that policy could reinforce (Supplementation file Table S1). Free-range systems dominated in dry (86%) and sub-humid (89%) zones, allowing behavioral thermoregulation through microhabitat selection. Intensive systems were more common in the humid zone (27%), potentially maladaptive under thermal stress unless combined with cooling infrastructure. Feeding practices (Figure 3) reflected local resource optimization: agricultural residues in the grain-producing dry zone (29%), fermented mash in the sub-humid zone (25%), and formulated feeds in the humid zone (29%). Stock selection was driven almost exclusively by phenotypic traits (98%), creating opportunities for participatory breed improvement programs that work within existing farmer decision frameworks.

**Figure 3.**
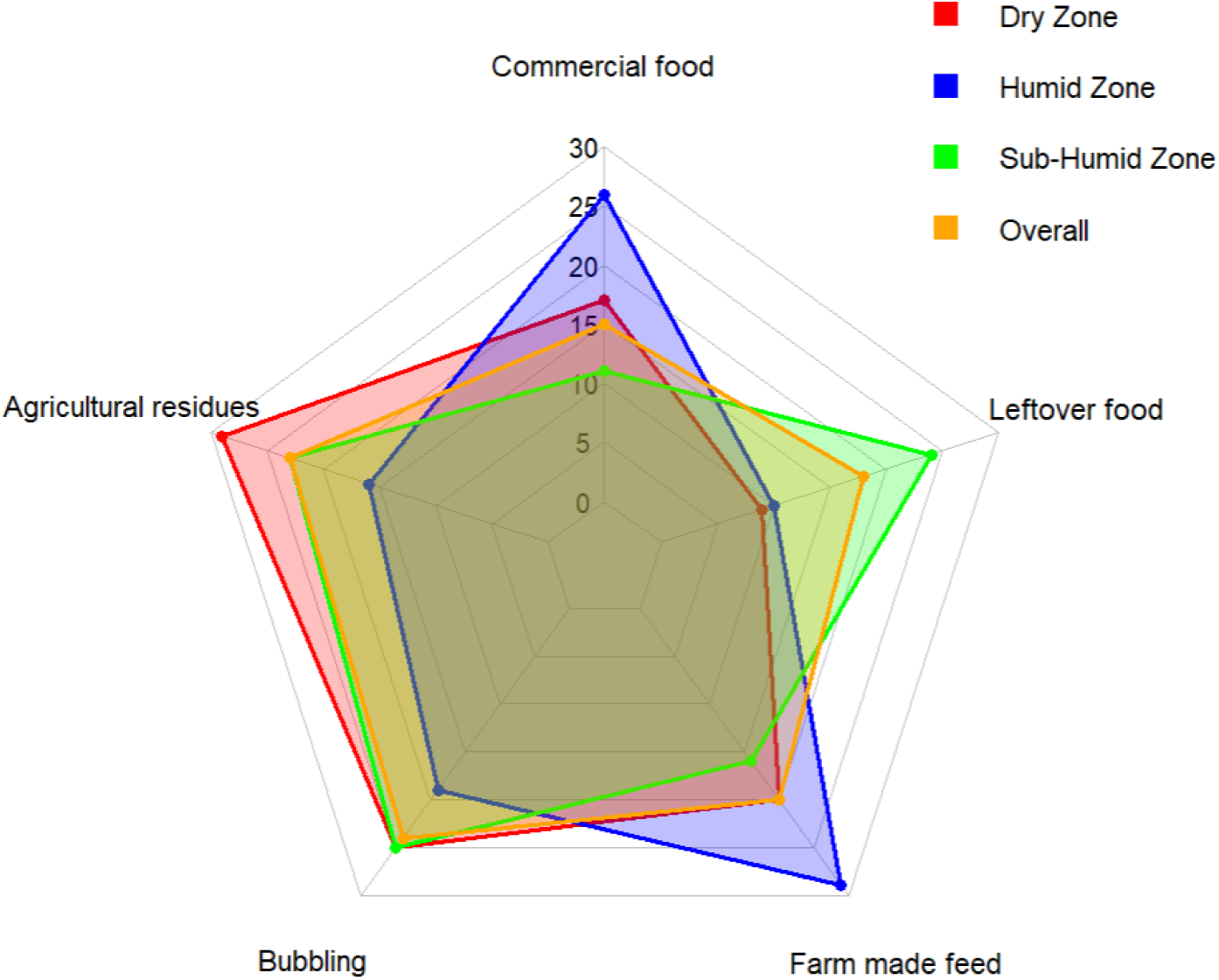
Radar chart illustrating the duck feed sources across three agroecological zones (Dry, humid, and sub-humid and the overall average). The five axes represent different feed types: Commercial food, Agricultural residues, Bubbling, Farm-made feed, and Leftover food. Agricultural residues refer to unprocessed by-products from crop cultivation, such as rice bran, maize husks, or groundnut shells. Bubbling denotes a locally prepared semi-fermented mash often based on maize or sorghum, mixed with water and occasionally enhanced with salt or other additives. Farm-made feed includes intentionally formulated mixtures using farm-available ingredients (e.g., grains, oilseed cakes, or kitchen by-products), typically blended in specific ratios to meet the birds’ dietary needs. Leftover food refers to cooked or uncooked human food remains provided to ducks, including starches, vegetable scraps, or soup residues.

### 3.3. Health Challenges: Disease Burden Across the Environmental Gradient

Health management patterns revealed zone-specific disease pressures and treatment gaps relevant to veterinary service planning (Figure 4 and Table 3).

**Figure 4.**
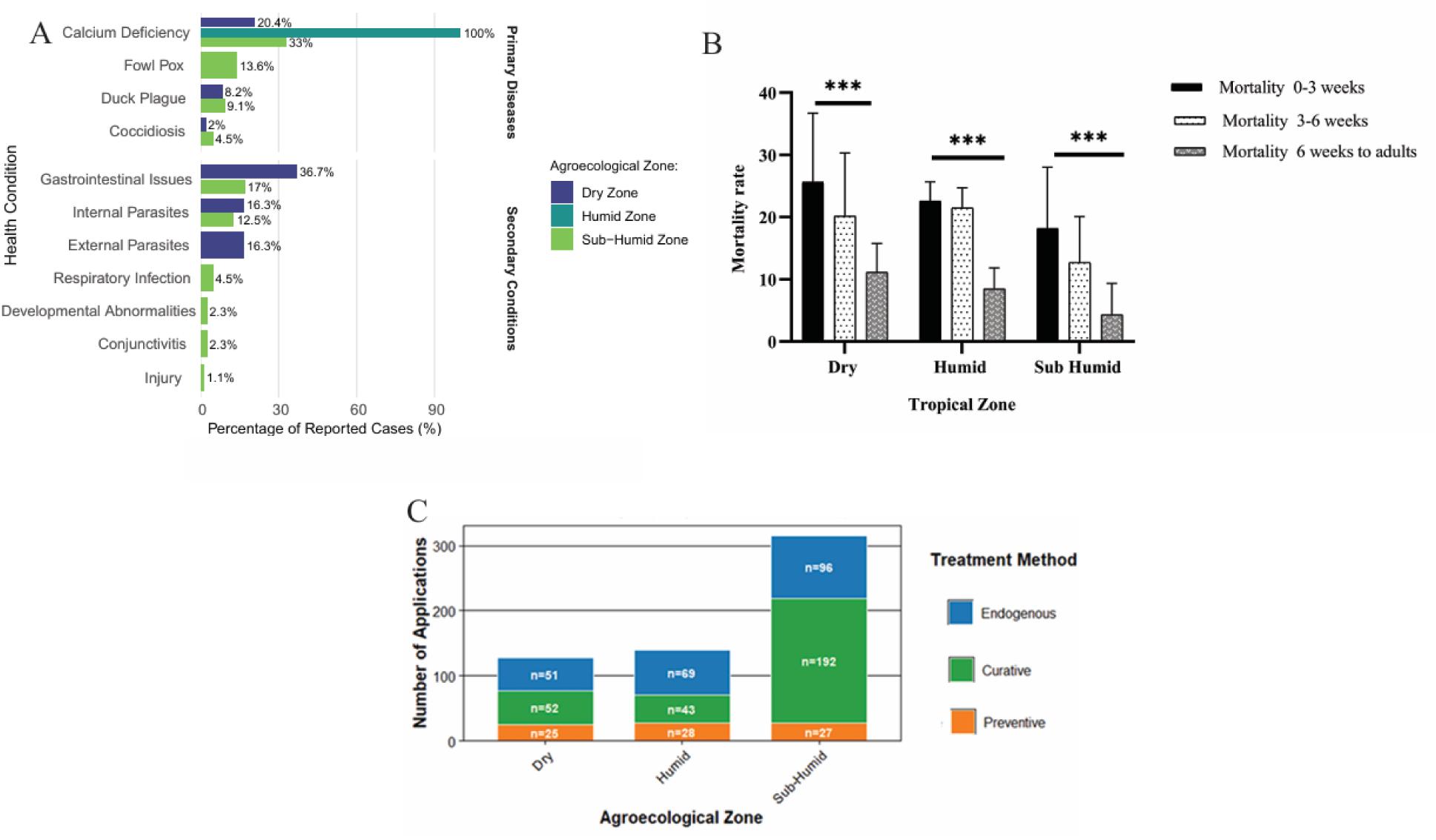
Distribution of disease prevalence, mortality, and therapeutic interventions across agroecological zones in ducks. The figure presents (A) the prevalence of primary diseases and secondary conditions by agroecological zone, including respiratory infections with influenza-like symptoms and gastrointestinal disorders such as diarrhea; (B) mortality rates stratified by agroecological zone and age group; and (C) the distribution of treatment methods across agroecological zones, shown as a stacked bar chart indicating the relative frequency of therapeutic approaches.

**Table 3:**
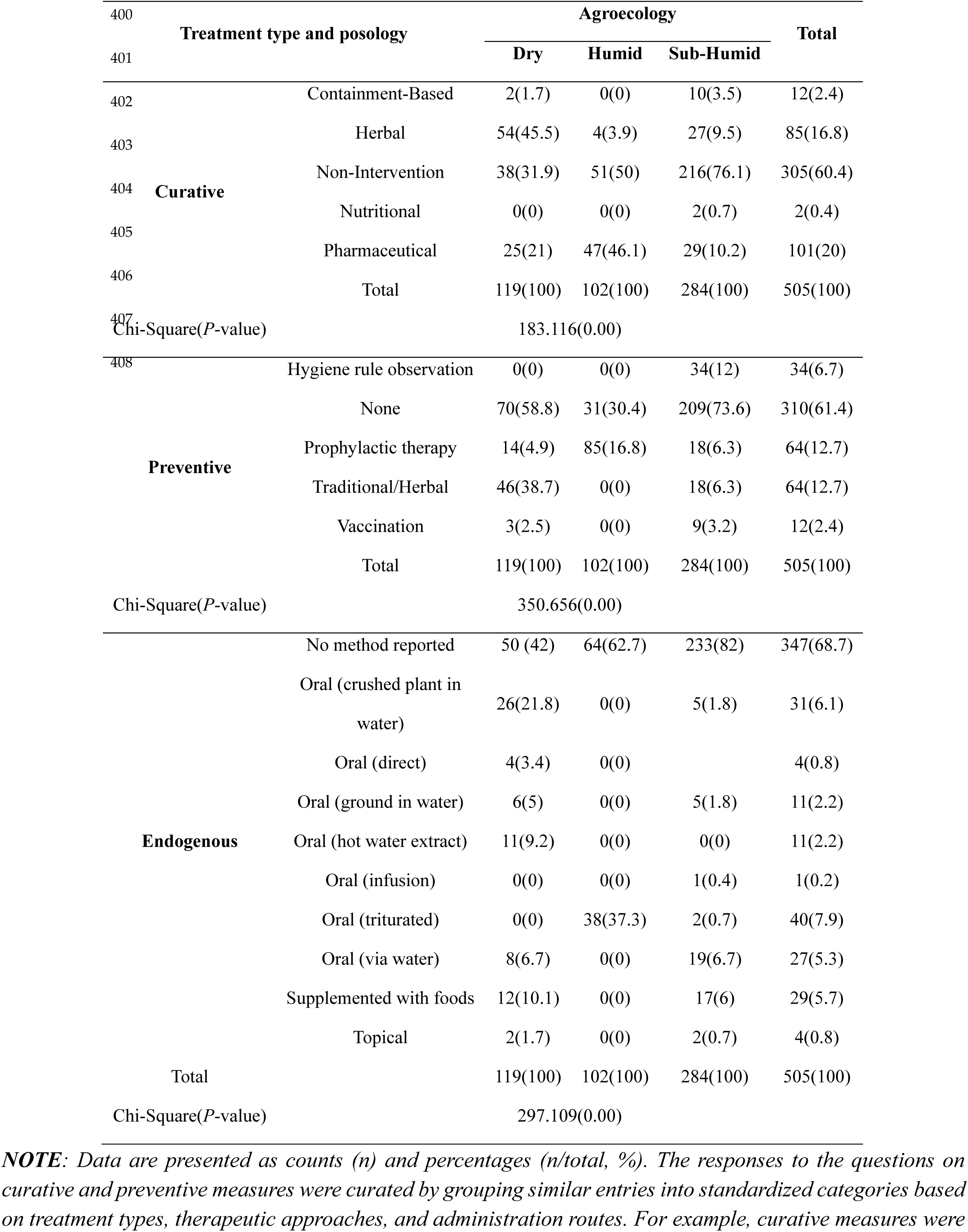
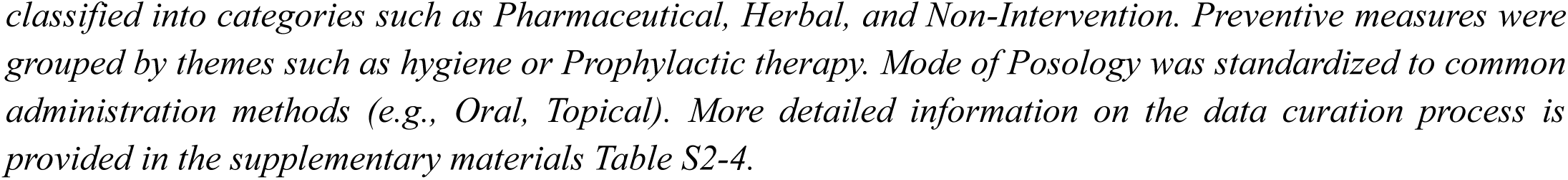
Descriptive statistics of curative, preventive, and posology measures across Agro-ecological zones.

Nutritional disorders predominated in the dry zone, with universal calcium deficiency indicating mineral imbalances in local feeds (Figure 4-A), addressable through extension programming on feed supplementation. Disease pressure intensified with humidity: the humid zone faced fowl pox (13.6%), duck plague (9.1%), and coccidiosis (4.5%), while the sub-humid zone experienced gastrointestinal disorders (36%) and parasitism (16%). These patterns suggest that climate-driven range shifts of pathogens may be occurring, consistent with FAO warnings about emerging disease risks under climate change.

Treatment practices reflected limited veterinary access (Figure 4-B). Endogenous therapies dominated in the humid (49.3%) and dry (39.8%) zones. Vaccination rates were negligible (less than 3.5%) across all zones despite the availability of effective vaccines for major duck diseases, representing a critical gap in disease prevention that policy intervention could address. Mortality patterns showed zone-specific temporal distributions: early mortality (0-3 weeks) peaked in the sub-humid zone (29.4%), while late mortality (greater than 6 weeks) was highest in the dry zone (10.7%), suggesting different selection pressures operating at different life stages (Figure 4-C).

### 3.4. Reproductive Performance: Quantifying Climate-Driven Productivity Losses

Reproductive performance data quantify the impact of climate impacts on productivity, providing baseline metrics for monitoring SDG 2 monitoring and for the economic assessment of adaptation benefits.

Egg production declined significantly along the thermal gradient (Figure 5-A). Lifetime production was highest in the dry zone (40.7 ±10.1 eggs), intermediate in the humid zone (38.4 ±5.06 eggs), and lowest in the sub-humid zone (37.1 ±9.4 eggs). Differentiation was most pronounced in third-cycle production: dry zone (17.0 ±4.12 eggs) versus sub-humid zone (14.2 ±3.83 eggs) (p = 1.57 × 10-10). Extrapolating these findings across Benin’s estimated duck population suggests substantial protein production losses in higher-stress zones.

**Figure 5.**
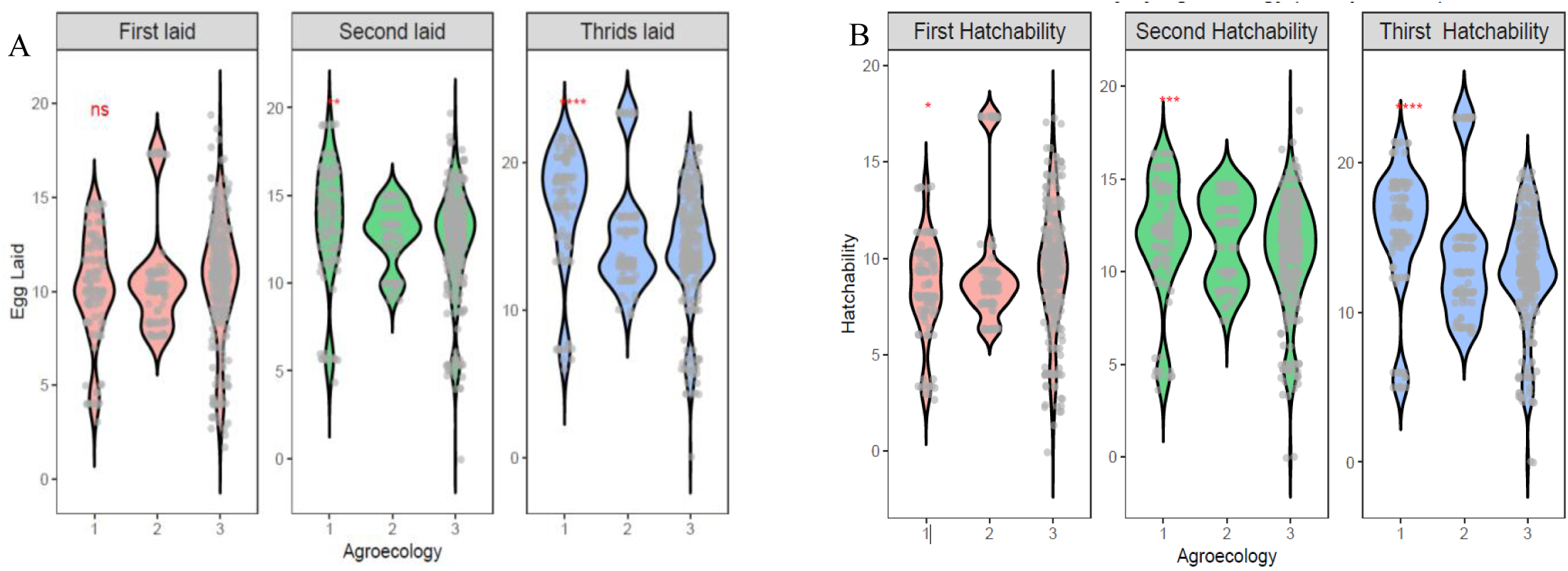
Egg production and hatchability performance across agroecological zones in Benin. The figure shows (A) the number of eggs laid and (B) hatchability rates across the dry tropical zone (1), humid tropical zone (2), and sub-humid tropical zone (3). Statistical significance is indicated as p < 0.05, p < 0.01, p < 0.001, and p < 0.0001.

Hatchability showed similar trends (Figure 5-B): dry zone (15.5 ± 4.16%), humid zone (13.7 ± 4.26%), and sub-humid zone (12.7 ± 3.68%) (p = 9.82 × 10-10). The 2.8 percentage point reduction between the dry and sub-humid zones translates to approximately 18% fewer ducklings reaching maturity per breeding cycle, representing a substantial constraint on flock replacement and expansion.

Phenotype-performance associations revealed locally adapted variants with superior fitness under native conditions, providing core evidence for breed-for-environment policies (Table 4). Black-and-white pied ducks were dominant in the dry (89%) and sub-humid (66%) zones and exhibited the highest hatchability (31% and 23%). White-black-green pied ducks were dominant in the humid zone (94%) and exhibited 32% hatchability under severe THI conditions. Beta regression confirmed that these associations remained significant after controlling for zone effects, indicating genuine phenotype-climate interactions rather than geographic confounding.

**Table 4:** Dominant phenotype (plumage color) frequency per agro-ecological zone.

| Dominant Phenotype | Tropical Zone |  |  |  |  |  |  |  | Pearson<br>Chi-Square | P-Value |
| --- | --- | --- | --- | --- | --- | --- | --- | --- | --- | --- |
|  | Dry |  | Subhumid |  | Humid |  | Overall |  |  |  |
|  | N | Percentage (%) | N | Percentage (%) | N | Percentage (%) | N | Percentage (%) |  |  |
| White, Green, and Blue Pied | 16 | 13.4 | 70 | 24.6 | 82 | 80.4 | 168 | 33.3 | 132.599 | 0.7e-7 |
| Black and White Pied | 106 | 89.1 | 188 | 66.2 | 18 | 17.6 | 312 | 61.8 | 124.035 | 0.000 |
| Brown and White Pied | 1 | 0.8 | 15 | 5.3 | 2 | 2.0 | 18 | 3.6 | 5.769 | 0.056 |
| Pearl-gray | 14 | 11.8 | 15 | 14.7 | 29 | 10.2 | 58 | 11.5 | 1.503 | 0.472 |
| white | 74 | 62.2 | 181 | 63.7 | 39 | 38.2 | 294 | 58.2 | 21.064 | 0.5e-4 |
| White and Green Pied | 6 | 5.0 | 41 | 14.4 | 4 | 3.9 | 51 | 10.1 | 13.524 | 0.1e-4 |
| black | 72 | 60.5 | 103 | 36.3 | 44 | 43.1 | 219 | 43.4 | 20.060 | 0.9e-5 |
| brown | 20 | 16.8 | 71 | 25.0 | 6 | 5.9 | 97 | 19.2 | 18.253 | 0.1e-5 |
| White, black, and green Pied | 38 | 31.9 | 123 | 43.3 | 96 | 94.1 | 257 | 50.9 | 99.905 | 0.6e-6 |

**NOTE:** Only affirmative responses (coded as "1") were considered in the analysis. Respondents were asked whether a given phenotype was present; responses indicating presence were coded as 1 and those indicating absence as 0. Frequency estimates reflect the distribution of the dominant phenotypes based solely on positive (affirmative) responses within each agroecological zone.

### 3.5 Morphometric Evidence for Thermoregulatory Adaptation

Principal component analysis provided multivariate evidence for climate-driven phenotypic differentiation (Figure 6). LDA (Figure 6-A) and PC1 (35.3% variance) represented body size; while PC2 (22.7% variance) captured limb and appendage proportions (Figure 6-B). Zone-specific clustering demonstrated non-random phenotypic distributions consistent with local adaptation, as expected if selection pressure varies systematically across the gradient (Figure 7 and Supplementation file Figure S2).

**Figure 6.**
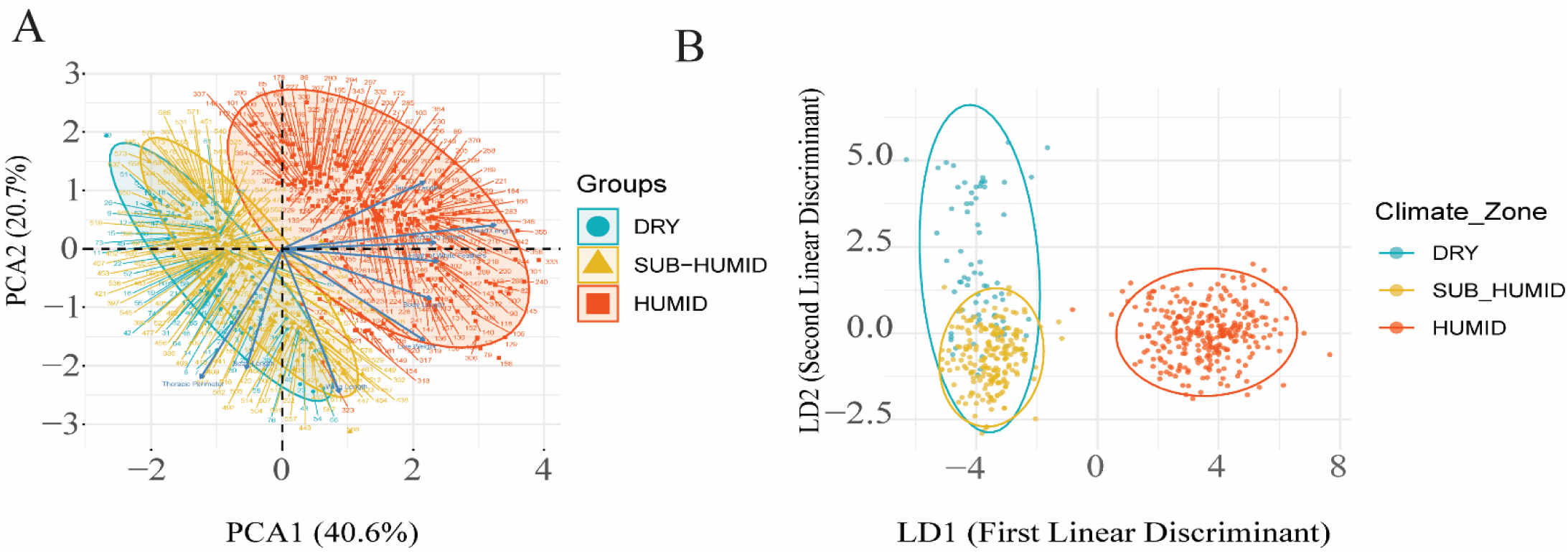
Multivariate analysis of morpho-biometric traits across climate zones. The figure illustrates (A) a principal component analysis (PCA) of key morpho-biometric traits by climate zone and (B) a linear discriminant analysis (LDA) showing the classification of individuals according to climate zone based on selected morpho-biometric traits.

**Figure 7.**
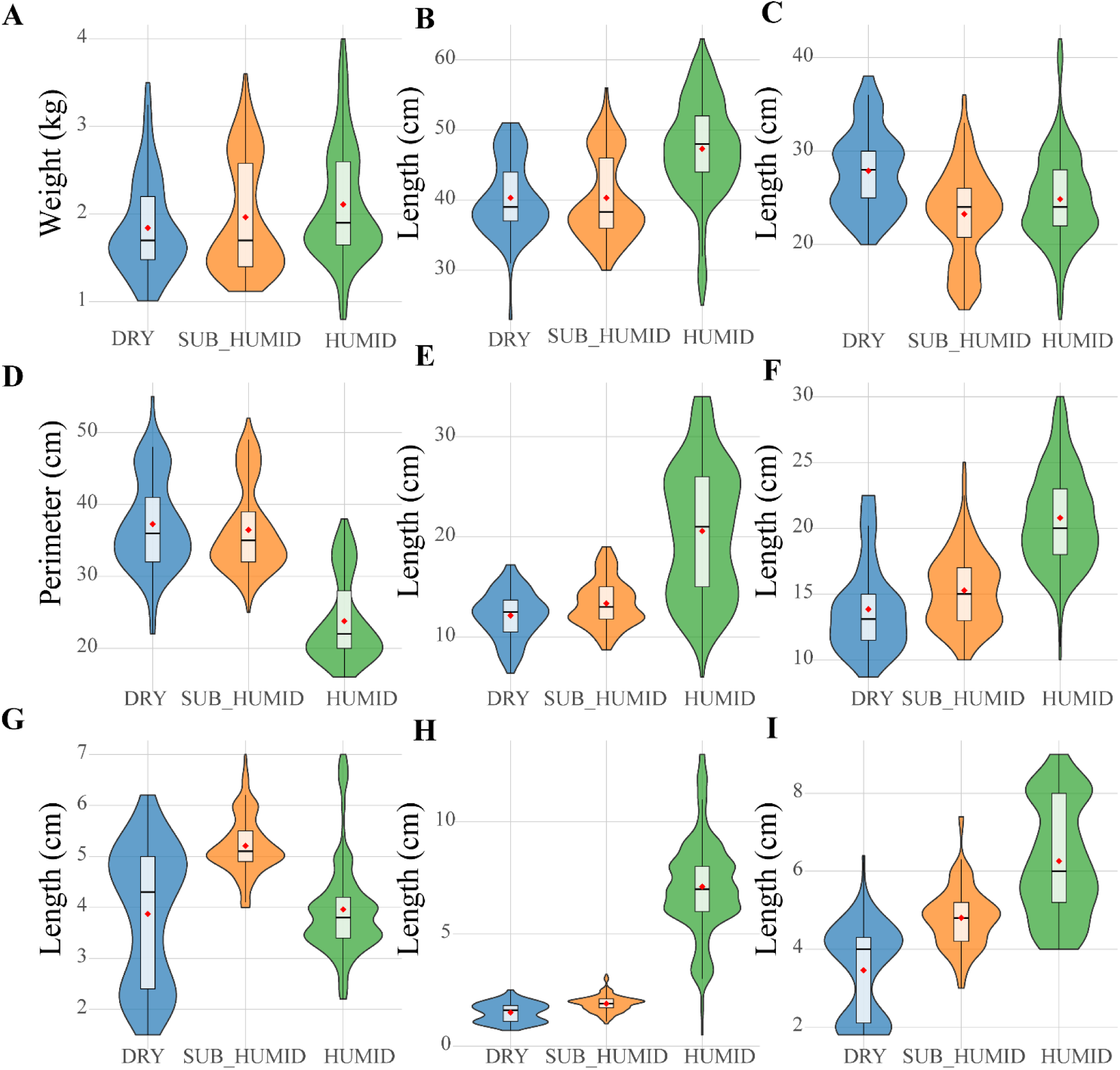
Morpho-biometric characteristics of ducks across different climate zones. The figure presents comparative analyses of (A) live weight, (B) body length, (C) wing length, (D) thoracic perimeter, (E) sternum length, (F) white feather length, (G) beak length, (H) tarsus length, and (I) head length across climate zones.

Thermoregulatory trait analysis revealed coordinated adaptation across multiple pathways (Table 5). Body volume-surface ratios were 42.7% lower in humid zones (p = 0.002), optimizing convective heat loss. Limb proportion ratios were 72.7% higher in humid zones (p ≤ 0.001), consistent with Allen’s rule predictions for enhanced heat dissipation through increased appendage surface area. White feather length was 57.5% greater (p ≤ 0.001) and white feather proportion 31.3% higher (p = 0.019) in humid zones, maximizing solar reflectance under high radiation conditions. These morphological patterns demonstrate that Benin’s Muscovy duck populations have differentiated along thermoregulatory axes in response to the climate gradient, providing empirical evidence that locally adapted genetic resources exist and could be strategically conserved and deployed under climate change scenarios.

**Table 5:** Spatial variation in thermoregulatory morphological traits of Muscovy ducks across Benin’s agro-ecological gradient.

| Thermoregulatory Trait | Dry Zone | Sub-humid Zone | Humid Zone | P-value |
| --- | --- | --- | --- | --- |
| Body Volume-Surface Ratio<br>(Thoracic Perimeter / Live Weight) | 21.49± 0.86 | 19.65± 1.13 | 12.32± 1.17 | 0.0017** |
| Beak Heat Dissipation Ratio<br>(Beak Length / Width) | 2.14± 0.07 | 2.46± 0.09 | 1.77± 0.09 | < 0.001*** |
| Limb Proportion Ratio<br>(Tarsus Length / Drumstick Length) | 0.39± 0.07 | 0.36±0.10 | 0.68± 0.10 | < 0.001*** |
| White Feather Length (cm) | 13.51± 0.56 | 15.27± 1.13 | 21.27± 1.17 | < 0.001*** |
| White Feather Proportion<br>(White Feathers / Body Length) | 0.35± 0.02 | 0.38± 0.006 | 0.46± 0.0054 | 0.0188* |

## 4. Discussion

### 4.1 Bridging the Science-Policy Gap: From Phenotypic Characterization to NDC Implementation

This study demonstrates how environmental assessment methodologies can generate policy-actionable evidence for integrating animal genetic resources into climate adaptation frameworks. The findings directly address three critical gaps limiting NDC implementation in the livestock sector: (1) a lack of quantitative data linking genetic resources to adaptation outcomes; (2) a lack of spatially explicit information for zone-specific policy targeting; and (3) an insufficient understanding of existing farmer adaptation practices that policy could reinforce. By providing this evidence base, the study offers a replicable model for other ECOWAS nations facing similar challenges in operationalizing Paris Agreement commitments.

Benin’s updated NDC identifies agriculture as a priority sector for climate adaptation and calls for the promotion of adapted varieties and breeds, but does not specify which breeds, which agro-ecological zones, or the empirical basis for such recommendations [28]. Our findings address this deficiency by demonstrating that black-and-white pied Muscovy ducks exhibit superior adaptation to dry and sub-humid zone conditions (31% and 23% hatchability respectively), while white-black-green pied variants maintain fitness under humid zone thermal stress (32% hatchability despite THI 78-82). These phenotype-zone associations provide the specificity required for implementing the NDC’s breed adaptation commitment through targeted conservation and dissemination programs.

The morphometric data also enable registration of Benin’s Muscovy duck populations in FAO’s Domestic Animal Diversity Information System (DAD-IS), fulfilling reporting obligations under the CBD Global Plan of Action for Animal Genetic Resources. Currently, no Beninese duck populations are registered in DAD-IS, a documentation gap that undermines national claims to genetic resource sovereignty and limits access to international support mechanisms for conservation [29]. The standardized characterization conducted in this study provides the data package required for breed registration and subsequent inclusion in national genetic resource conservation strategies.

### 4.2 Climate Vulnerability Beyond IPCC Projections: Implications for Regional Planning

The climate analysis revealed warming trends (+0.31 to +0.38 degrees C per decade) exceeding IPCC AR6 median projections for West Africa (+0.25 degrees C per decade under SSP2-4.5) [30]. This finding has significant implications for regional adaptation planning under ECOWAS frameworks. If current trends continue, the sub-humid zone, home to the largest share of Benin’s duck farmers (56% of our sample), will transition from chronic moderate stress (THI 74-78) to severe stress conditions (THI greater than 78) within 15-20 years. This transition threshold corresponds to the stress regime currently experienced in the humid zone, where our data document 8.8% lower lifetime egg production and 18% lower duckling survival compared to the dry zone.

These projections are directly relevant to the ECOWAS Climate-Smart Agriculture investment framework, which emphasizes no-regret adaptation strategies that yield benefits across plausible climate scenarios [31]. Our results indicate that anticipatory deployment of humid-zone-adapted phenotypes represents such a no-regret intervention, consistent with resilience-building principles articulated under the Sendai Framework for Disaster Risk Reduction [32]. Accelerated warming in humid coastal zones further challenges assumptions that these regions will function as long-term climate refugia. Recent assessments caution that refugia capacity may erode when warming rates exceed historical variability and teh physiological tolerance limits of agricultural systems [30], underscoring the need to incorporate livestock-specific thermal thresholds into Benin’s National Adaptation Plan and ECOWAS regional climate modeling efforts.

### 4.3 Thermoregulatory adaptation and empirical support for FAO’s breed-for-environment paradigm

The observed morphometric differentiation across Benin’s environmental gradient provides direct empirical support for the FAO’s “breed-for-environment” paradigm, as advocated by Ouda and Zohry (33). This paradigm emphasizes matching animal genetic resources to local climatic constraints rather than relying on uniform genetic improvement through exotic germplasm. While this principle has been widely articulated in climate-smart livestock frameworks, quantitative evidence demonstrating its operation in smallholder poultry systems remains scarce in West Africa. Our results show that Beninese Muscovy duck populations have already undergone coordinated thermoregulatory differentiation, yielding phenotypes that outperform alternative variants under native thermal regimes, thereby validating the practical relevance of this approach.

The marked increase in limb proportion ratios in humid-zone ducks aligns with the thermoregulatory logic of Allen’s rule, where larger appendages can increase dry heat dissipation capacity under chronic heat load. Recent comparative analyses and mechanistic work in birds reinforce that appendage dimensions are tightly linked to thermal adaptation and heat-loss regulation [25]. Simultaneously, the heat-stress literature shows that sustained high temperature-humidity exposure depresses poultry productivity and survival, underscoring why selection on thermoregulatory morphology can have direct fitness consequences under tropical conditions [34].

Plumage traits displayed equally strong signatures of thermal adaptation. Ducks originating from humid zones exhibited significantly greater white feather length and higher proportions of white plumage, traits associated with reduced solar heat gain through enhanced reflectance and lower feather surface temperatures under high radiation loads. Recent experimental and comparative studies in poultry and wild birds confirm that lighter plumage reduces radiant heat absorption and mitigates heat stress, contributing to improved thermoregulation and performance under hot, humid conditions [35, 36]. These effects are particularly pronounced under chronic thermal stress, where radiative heat load becomes a dominant constraint on productivity.

Unlike physiological or genomic indicators that require laboratory infrastructure, plumage coloration constitutes a visually accessible and farmer-recognizable proxy for thermal adaptation. Contemporary poultry studies demonstrate that light-colored birds consistently maintain lower body temperatures, improved survivability, and more stable reproductive performance under heat stress compared with dark-plumaged counterparts [37]. This makes plumage-based selection especially relevant for smallholder systems where climate adaptation must rely on low-cost, observational criteria.

Taken together, the coordinated differentiation of limb proportions and plumage traits in Muscovy ducks conforms to established ecogeographic principles and yields operationally simple selection markers for climate-resilient breeding. By translating the FAO’s “breed-for-environment” guidance into measurable, zone-specific phenotypic indicators, these findings demonstrate how animal genetic resource management can be integrated into climate adaptation policies without dependence on advanced genomic infrastructure.

### 4.4. Productivity losses under climate stress and implications for food security investment

The reproductive performance data allow for the quantification of climate-attributable productivity losses. This fills an evidence gap that has limited cost-benefit assessments of livestock adaptation options within CAADP-aligned investment planning. The observed reduction in lifetime egg production between dry and sub-humid zones (40.7 vs. 37.1 eggs per female) corresponds to an 8.8% decline in reproductive output associated with elevated thermal stress. Combined with the 18% reduction in duckling survival, the total effective reproductive yield per breeding female is approximately 25% lower in higher-THI environments. This is consistent with documented heat-stress effects on poultry reproduction in tropical systems [37].

Scaling these differentials to the national level demonstrates their policy relevance. With Benin’s duck population estimated at 0.5-0.7 million birds and breeding females comprising roughly 60% of that population [38], productivity disparities between lower-and higher-stress zones plausibly translate into substantial annual losses in egg production and surviving ducklings. While such extrapolations necessarily involve uncertainty, similar order-of-magnitude assessments have been used to inform climate-risk screening and investment prioritization in smallholder livestock systems [31, 39]. Under prevailing market conditions, reduced duckling survival alone represents a non-trivial loss of household income, reinforcing the vulnerability of poultry-dependent livelihoods to incremental warming.

Importantly, these estimates support a positive economic case for anticipatory adaptation. Partial recovery of climate-related losses through targeted measures such as deployment of heat-adapted phenotypes, improved incubation practices, or low-cost housing modifications would likely generate benefits exceeding typical intervention costs. This type of quantified productivity evidence aligns with the Green Climate Fund’s emphasis on adaptation actions with demonstrable livelihood impacts and supports the inclusion of indigenous poultry systems within climate finance and national adaptation planning.

### 4.5 Production systems as indigenous adaptation and targets for extension policy

The zone-specific management patterns observed in our survey are consistent with behavioral thermoregulation strategies documented in poultry under hot climates. The high reliance on free-range management in drier zones likely enables birds to reduce heat load by seeking shade, exploiting airflow, and shifting activity to cooler periods, compared with continuous confinement this can partially buffer productivity under thermal stress [40]. Rather than treating extensive systems as merely “traditional,” extension policy can frame them as context-adapted risk management. Policy should focus on improving low-cost components that enhance heat avoidance (shade access, water placement, resting sites) while preserving mobility-based coping capacity [41].

Feeding differentiation across zones similarly reflects farmer optimization under local resource constraints. Fermented feeds have growing empirical support in poultry for improving nutrient availability, gut microbial balance, and performance. Recent studies specifically demonstrate benefits in ducks, including Muscovy ducks, under practical feeding conditions [42]. In this context, the prominence of fermented mash in the sub-humid zone is plausibly adaptive and should be evaluated through participatory trials that quantify performance and cost trade-offs rather than replaced by uniform “improved feeding” packages [42].

Finally, the near-universal reliance on phenotypic selection highlights a realistic entry point for genetic improvement: farmer selection commonly targets survival and reproductive success under local stressors. Evidence from smallholder livestock and village poultry systems shows that farmer-defined trait priorities can be integrated into community-based or participatory breeding approaches to deliver locally acceptable genetic gains while maintaining adaptive diversity

### 4.6 Health–climate interactions and emerging veterinary preparedness needs

The spatial concentration of viral disease reports in humid zones is consistent with broader evidence indicating that climate variables especially humidity and rainfall patterns can alter pathogen persistence and vector ecology. This in turn increase transmission opportunities and shift disease risk across landscapes [43]. For avian pox, public health and veterinary guidance highlights its higher occurrence in warm, humid areas closely linked with strong links to mosquito cycles. This supports the plausibility of humidity-associated clustering and potential expansion under wetter/warmer conditions [44]. Regarding duck plague (duck virus enteritis), authoritative veterinary standards emphasize its high contagion and management relevance for ducks, underscoring the need for preparedness in the areas where duck production is intensifying [45].

The very low vaccination uptake observed in our sample therefore represents a manageable policy bottleneck: where vaccines and delivery mechanisms exist, adoption commonly depends on logistics, cost, and knowledge transfer; classic extension constraints. In parallel, the concentration of early mortality (0–3 weeks) in the sub-humid zone is consistent with the vulnerability of young birds to combined thermal and disease stress, indicating that targeted extension packages focused on brooding microclimate (shade/ventilation), water access, and hygiene could deliver rapid gains without requiring genetic change [46].

### 4.7 Policy Recommendations: From Evidence to Action

Based on these findings, we recommend the following policy actions at national, regional, and international levels:

**At the national level, Benin should:**

(1) incorporate the phenotype-zone associations identified in this study into the National Livestock Development Strategy currently under revision, specifying black-and-white pied Muscovy ducks for dry and sub-humid zone promotion and white-black-green pied variants for humid zone programs;
(2) Register characterized Muscovy duck populations in FAO’s DAD-IS database to fulfill CBD reporting obligations and enable access to international genetic resource conservation mechanisms; (3) Establish a national Muscovy duck breeding nucleus with zone-adapted populations for germplasm conservation and farmer access;
(4) Integrate duck health programming into the national veterinary service mandate, including vaccination campaign protocols and extension messaging on disease prevention;
(5) include livestock thermal stress monitoring in the national climate early warning system managed by the Agence Nationale de la Meteorologie.

**At the ECOWAS regional level:**

(1) Incorporate animal genetic resource conservation into the ECOWAP Climate-Smart Agriculture investment framework, using this study’s methodology as a model for replication across member states;
(2) Establish regional gene banks for climate-adapted poultry genetic resources, enabling cross-border germplasm exchange when local populations face climate threats;
(3) Integrate livestock productivity monitoring into regional climate vulnerability assessment protocols.

At the international level:

(1) Strengthen reporting requirements under the CBD Global Plan of Action for Animal Genetic Resources to include climate adaptation indicators;
(2) Establish Green Climate Fund financing windows for animal genetic resource conservation as climate adaptation;
(3) Incorporate livestock breed-environment matching into NDC implementation guidance provided by UNFCCC technical support mechanisms.

### 4.8 Study limitations and research priorities

The cross-sectional design captures phenotypic variation at a single time point but cannot directly demonstrate evolutionary change; longitudinal monitoring would strengthen causal inference. Priority research needs include: (1) genomic analysis to identify the genetic basis of thermoregulatory traits, enabling marker-assisted selection; (2) physiological studies characterizing heat tolerance mechanisms under controlled conditions; (3) economic analysis of adaptation intervention cost-effectiveness; (4) longitudinal monitoring to detect phenotypic change under continued warming; and (5) replication across ECOWAS countries to assess regional generalizability. These research priorities align with CAADP’s Pillar IV mandate for agricultural research supporting continental development goals.

## 5. Conclusion

This study provides the first comprehensive evidence base for integrating Muscovy duck genetic resources into climate adaptation policy in West Africa. By demonstrating coordinated thermoregulatory adaptation across Benin’s environmental gradient, including morphological specialization consistent with ecogeographic principles, phenotype-specific reproductive performance under different thermal stress regimes, and climate-responsive production systems, the findings establish that locally adapted genetic resources exist and can be strategically managed for climate resilience.

The policy implications are substantial and actionable. The phenotype-zone associations identified provide the specificity required for implementing Benin’s NDC commitment to promote adapted varieties and breeds in the livestock sector. The morphometric data enable fulfillment of CBD reporting obligations through DAD-IS registration. The productivity loss quantification demonstrates the economic case for adaptation investment in GCF funding proposals. The production system characterization identifies leverage points for extension programming that reinforces rather than replaces indigenous adaptation practices.

More broadly, this study demonstrates how environmental assessment methodologies can bridge the persistent gap between international biodiversity commitments and national policy implementation. The approach, systematic phenotypic characterization across environmental gradients, quantification of performance under different stress regimes, and integration of socioeconomic data on management practices, is replicable across species and regions facing analogous challenges. As climate change intensifies pressure on agricultural systems throughout sub-Saharan Africa, such evidence-based approaches to genetic resource management will be essential for achieving both the Paris Agreement’s adaptation goals and the Sustainable Development Goals’ food security targets.

The Muscovy duck populations of Benin exemplify a broader reality: centuries of farmer selection under heterogeneous environmental conditions have produced locally adapted genetic resources that represent both products of past adaptation and tools for future resilience. Recognizing, conserving, and strategically deploying this adaptive diversity is not merely a technical exercise in animal breeding; it is a core component of climate-resilient development policy that determines the capacity of livestock-dependent communities to sustain their livelihoods under an uncertain climate future.

## Supporting information

**Table S1:** Breeding methods and habitat types across tropical zones in duck populations.

**Table S2:** Duck Feed index for each zone

**Table S3:** Categorization of Curative Measures Reported by Respondents

**Table S4:** Curated categories Methods.

**Table S5:** Classification Key for Harmonizing Modes of Posology

**Figure S1:** Breeding objectives and mating systems of ducks across tropical zones.

**Figure S2:** Model-based hatchability predictions incorporating zone-specific thermal-humidity index, plumage phenotype, and agroecological zone field data.

**SE1**: Methodology for Curation and Categorization of Curative Measures

## Notes

### Competing Interest Statement

The authors have declared no competing interest.

### Funding

- MOF and MARA: CARS-42-2
- IFS: I3-6655-1

